# A MTA2-SATB2 chromatin complex restrains colonic plasticity toward small intestine by retaining HNF4A at colonic chromatin

**DOI:** 10.1101/2022.12.15.520587

**Authors:** Wei Gu, Xiaofeng Huang, Pratik N. P. Singh, Ying Lan, Jesus M Gomez-Salinero, Shahin Rafii, Mike Verzi, Ramesh Shivdasani, Qiao Zhou

## Abstract

Plasticity between cell lineages is a fundamental but poorly understood property of regenerative tissues. In the gut tube, small intestine absorbs nutrients whereas colon absorbs electrolytes. In a striking display of inherent plasticity, adult colonic mucosa lacking the chromatin factor SATB2 is converted to small intestine. Using proteomics and CRISPR-Cas9 screen, we identified MTA2 as a crucial component of the molecular machinery that, together with SATB2, restrain colonic plasticity. MTA2 loss in adult mouse colon activated lipid absorptive genes and functional lipid uptake. Mechanistically, MTA2 co-binds with HNF4A, an activating pan-intestine transcription factor (TF), on colonic chromatin. MTA2 loss leads to HNF4A release from colonic and gain on small intestinal chromatin. SATB2 similarly restrains colonic plasticity through a HNF4A-dependent mechanism. Our study provides a generalizable model of lineage plasticity in which broadly-expressed TFs are retained on tissue-specific enhancers to maintain cell identity and prevent activation of alternative lineages; their release unleashes plasticity.

## Introduction

Cells in regenerative tissues can exhibit substantial phenotypic plasticity upon injury^1-3^. Differentiated quiescent cells may dedifferentiate along its lineage trajectory to become progenitors or stem cells, then assume forward differentiation to produce progenies for tissue repair, as reported in the lung, the intestine, and the skin^4-7^. Cells could also cross lineage boundaries and switch fate. Examples of lineage plasticity include conversion of hepatocytes to cholangiocytes in the liver, alpha/delta to beta cells in the pancreas, and hair follicle stem cells to epidermal stem cells in the skin^8-12^. Lineage plasticity must be tightly regulated because undesirable plastic events, such as metaplasia, promote dysfunction and could serve as precursors to tumorigenesis^13,14^. Some of the signaling pathways, transcriptional mediators, and chromatin substrates of cellular plasticity have begun to emerge from recent studies^8,15-18^. Nevertheless, our understanding of the molecular complexes that safeguard cellular identity and mediate lineage plasticity remains limited.

The small and large intestines are markedly different in cell composition and function^19^. While the small intestine absorbs nutrients through the enterocytes that line the mucosal surface, colon mainly absorbs electrolytes and water. Accordingly, enterocytes express many transporters for lipids, amino acids and carbohydrates that the colonocytes lack^20^. Nevertheless, our genome-wide mapping with the enhancer maker H3K4me1 and with ATAC-seq (Transposase Accessible Chromatin with high throughput sequencing) showed that colonic epithelial cells surprisingly harbor primed small intestine enhancers, suggesting an intrinsic permissiveness for small intestine gene activation in colon^21^. Upregulation of small intestine genes has been reported in the colon of patients with short bowel disease^22^ whereas a more substantial, colon to small intestine transcriptomic shift may occur in some patients with inflammatory bowel diseases (IBD)^23^. Colon thus displays various degrees of cellular plasticity.

We recently identified SATB2, a colon-restricted chromatin factor, as a central regulator of colonic gene expression and lineage plasticity^21^. *Satb2* deletion converts colonic stem cells into small intestine ileum-like stem cells, leading to the replacement of colonic mucosa by ileum-like mucosa in adult mouse colon. SATB2 is also expressed in mature colonocytes, where loss of SATB2 rapidly activates ileal genes. Despite these observations, mechanisms of colonic plasticity remain largely unknown.

We hypothesized that some of the SATB2-associcted chromatin factors may regulate colonic plasticity. Using Affinity Purification and Mass Spectrometry (AP-MS), we identified SATB2-associated proteins in colonic epithelium, including MTA2 (metastasis associated protein 2) and multiple members of the NuRD (Nucleosome Remodeling Deacetylase) complex. CRISPR (clustered regularly interspaced short palindromic repeats)-mediated functional evaluation in colonic organoids suggest that MTA2-containing NuRD complex could regulate colonic transcription. MTA2 expression is restricted to colonocytes and *Mta2* deletion in adult mouse colon activated small intestine genes, including fat absorptive genes, enabling lipid uptake by colonocytes. Mechanistic studies indicate that MTA2 and NuRD did not directly bind and silence small intestine genes, rather, MTA2 and HNF4A, a crucial intestinal TF, extensively co-occupied colonic chromatin. MTA2 loss led to HNF4A depletion at colonic and gain at small intestine enhancers, indicating a critical function for MTA2 in retaining HNF4A at colonic enhancers. Moreover, MTA2 physically interact with SATB2 and both restrain HNF4A on colonic chromatin, albeit at different strength, leading to different degrees of plasticity in *Mta2* vs *Sat2* mutant colon. Our data revealed a MTA2-SATB2 chromatin complex at colonic enhancers that perform the dual functions of safeguarding transcriptional fidelity and regulating plasticity. By restraining HNF4A, a TF expressed in both small and large intestines, at colonic enhancers, small intestinal gene transcription is blocked in colon; release of this block unleashes plasticity.

## Results

### MTA2-containing NuRD associates with SATB2 and regulates colonic gene expression

SATB2 and its homolog SATB1 have been proposed as chromatin hubs that orchestrate protein-protein and protein-DNA interactions^24,25^. Reasoning that some of the SATB2-associated factors may regulate colonic plasticity, we purified protein complexes that contain SATB2 from murine colonic glands (Fig. 1A and Fig. S1A). Two independent AP-MS experiments identified a total of 628 proteins with a false discovery rate (FDR) < 1%. Of these, 78 proteins were significantly enriched in both samples (SATB2 AP-MS signal intensity and MS count over IgG controls > 2 fold), with SATB2 itself being the most enriched (Fig. 1B).

**Figure 1.**
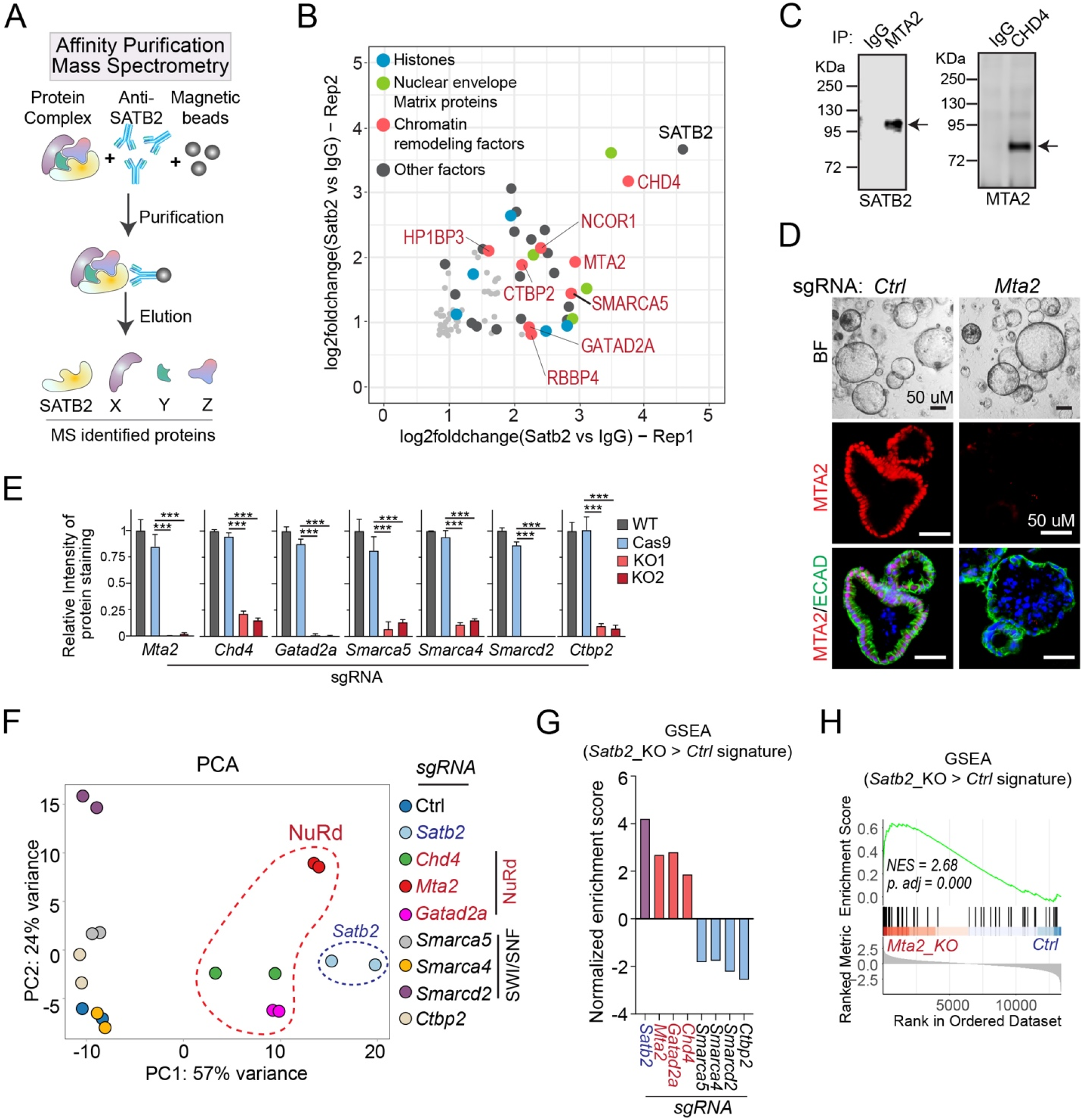
MTA2 and NuRD complex associate with SATB2 and regulate colonic transcription. **(A)** Candidate SATB2-associated proteins were identified from mouse colonic glands by affinity purification (AP) with anti-SATB2 antibody followed by Mass Spectrometry (MS). **(B)** Of the 68 proteins enriched in both AP-MS experiments, the top 40 (highlighted as colored balls) contained many histones, matrix proteins and chromatin remodeling factors. **(C)** Co-IP demonstrated interactions of SATB2 with MTA2 and MTA2 with CHD4, a core member of NuRD complex. **(D)** CRISPR-CAS9 and gRNA was used to successfully disrupt *Mta2* from cultured mouse colonic organoids, as shown by immunofluorescence staining. BF: bright field. ECAD: E-cadherin. **(E)** Immunoblot quantification showed significant reduction of seven SATB2-associated chromatin factors after CRISPR-mediated deletion in colonic organoids. Two independent CRISPR experiments and two controls were shown. Mean ± S.D. *** P < 0.001. P value by unpaired t-test. (**F-H**) RNA-sequencing showed that disrupting NuRD members (*Chd4, Mta2* and *Gatad2a*) but not SWI/SNF factors (*Smarca5, Smarca4* and *Smarcd2*) or *Ctbp2* caused transcriptomic shifts of colonic organoids toward that of *Satb2* knockout, illustrated by Principal Component Analysis (PCA, **F**) and Gene Set Enrichment Analysis (GSEA, **G, H**). NES: normalized enrichment score.

The top 40 candidate SATB2-associated proteins included an abundance of histones, nuclear matrix proteins, and chromatin remodeling factors, consistent with the proposed role of SATB family as chromatin organizers (Fig. 1B and Fig. S1B). Four members of the NuRD complex, including CHD4, MTA2, RBBP4 and GATAD2A, were among the top 40 interactors, suggesting association of NuRD complex with SATB2 (Fig. 1B). Using co-immunoprecipitation (co-IP), we observed interaction of SATB2 with MTA2 and of MTA2 with the NuRD core subunit CHD4 (Fig. 1C and Fig. S1C). We also validated interaction of SATB2 with SMARCD2 and SMARCA4, two members of the SWI/SNF chromatin remodeling complex identified in AP-MS (Fig. S1C).

To evaluate the functional importance of candidate SATB2-associated factors in colonic transcription, we used CRISPR-CAS9 to disrupt in murine colonic organoids nine chromatin remodeling genes whose protein products were enriched in our AP-MS analysis (Fig. S1D); of these, seven achieved deletion efficiencies of 80-95% by immunoblot analysis (Fig. 1D, 1E and Fig. S1E). RNA-sequencing indicated that deletion of *Chd4, Mta2*, or *Gatad2a*, but not the other factors, significantly altered colonic transcriptomes toward that of *Satb2* knockout organoids (Fig. 1F-H and Fig. S1F and 1G). These data suggest that the NuRD complex interacts with SATB2 and is functionally important in regulating colonic transcription.

### Activation of lipid absorptive genes in MTA2-deficient colonocytes

The colonic mucosa is a regenerative epithelium, with 4- to 7-day cycles of self-renewal powered by LGR5^+^ intestinal stem cells (ISCs) in the crypts of Lieberkuhn^26^. The colonic ISCs produce progenitors (transient amplifying cells) which give rise predominantly to absorptive colonocytes and secretory goblet cells (Fig. 2A). Immunohistochemistry revealed prominent MTA2 expression in upper, but not lower colonic glands, and in scattered sub-epithelial cells (Fig. 2B, 2C and Fig. S2A). In contrast, the NuRD subunits CHD4 and GATAD2A were present throughout the colonic epithelium, similar to SATB2 (Fig. 2B). The majority of MTA2^+^ cells (68.0 ± 7.6%) were CA1^+^ colonocytes and conversely, nearly all CA1^+^ mature colonocyes were MTA2^+^ (Fig. 2C). A minority of MTA2^+^ cells were goblet cells (6.5 ± 3.1%), recognized by Alcian blue stain (Fig. 2D-G). LGR5^+^ colonic stem cells did not express MTA2 (Fig. 2E and 2G). Thus, a MTA2-containing NuRD complex is enriched in terminally differentiated colonocytes on the luminal surface.

**Figure 2.**
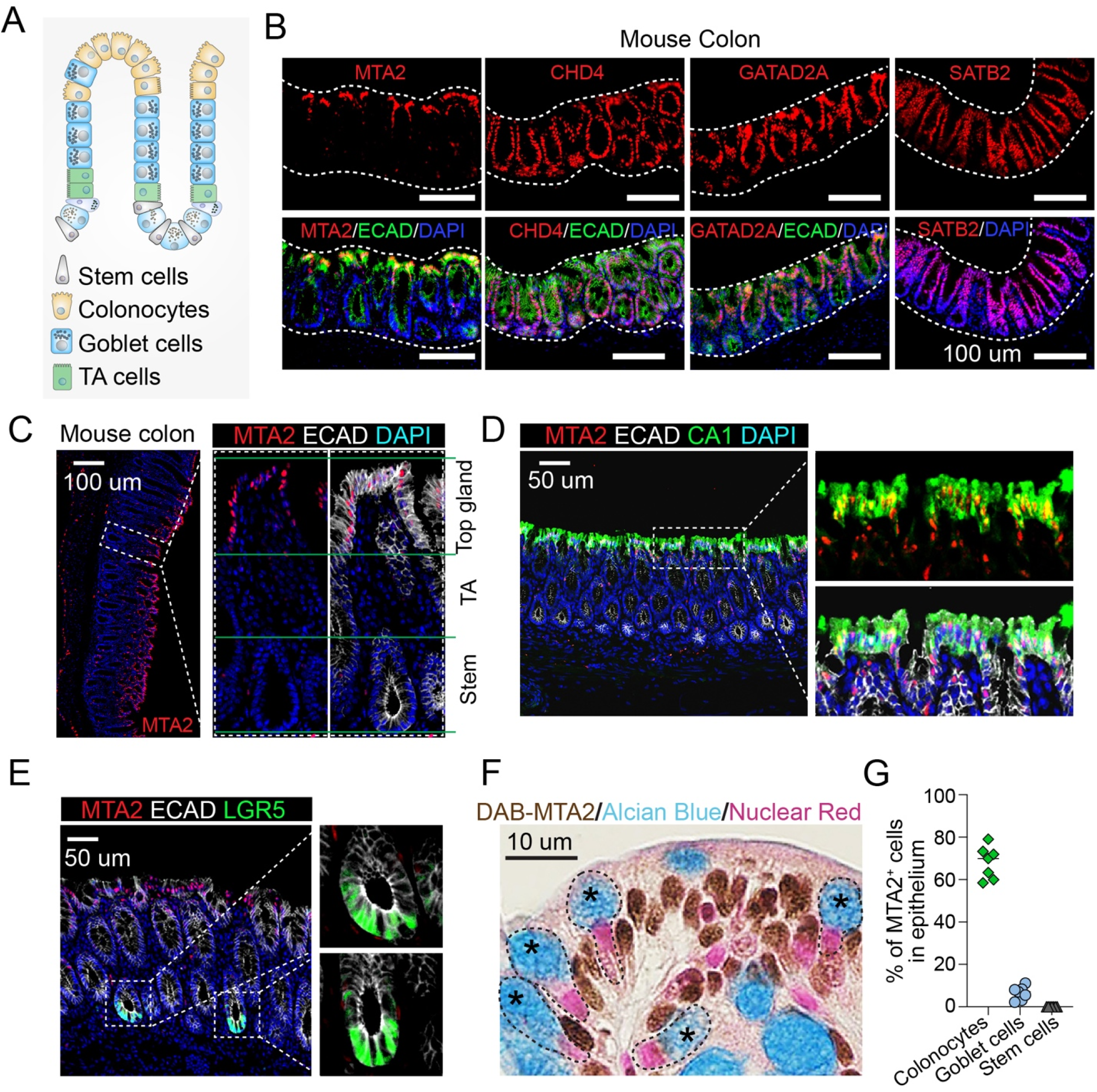
MTA2 expression is enriched in colonocytes. **(A)** Diagram of colonic epithelium and the cell lineages. Mature colonocytes are concentrated in the upper glands of mucosal surface. TA cells: transient amplifying cells. **(B)** Immunofluorescence staining showed prominent MTA2 expression in the upper glands of adult mouse colon whereas SATB2 and two other members of NuRD, CHD4 and GATAD2A, were present throughout the epithelium. (**C-E**) MTA2 was expressed in Ecad^+^CA1^+^ mature colonocytes (**C, D**), but not in LGR5^+^ colonic stem cells €. **(F)** Histology stain for MTA2 and Alcian Blue stain for goblet cells showed few goblet cells expressing MTA2 (**F**). **(G)** MTA2 was present mostly in colonocytes. N = 7 independent samples.

*Mta2* gene deletion from the intestinal mucosa of 2-month old *Villin-Cre*^*ER*^; *Mta2*^*fl/fl*^ mice (Fig. 3A, hereafter referred to as Mta2^cKO^) led to near complete absence of MTA2 (Fig. 3B). One month after Tamoxifen treatment, Mta2^cKO^ mice showed no overt changes in colonic histology or cell proliferation (Fig. S3A). RNA-sequencing of colonic glands revealed 200 up-regulated and 68 down-regulated genes (log_2_ fold change [LFC] > 1, adjusted p [padj] < 0.05) (Fig. 3C) in Mta2^cKO^ vs control colon. Mta2^cKO^ colon was enriched for functional pathways and gene sets in fat digestion and absorption, thiamine metabolism and chemokine signaling (Fig. 3D) (Fig. 3E, 3F). Transporters for amino acids, carbohydrates, bile salt and vitamins also exhibited a trend of up-regulation in Mta2^cKO^ colon (Fig. S3B). In contrast, no pathway was significantly enriched among the down-regulated genes (P < 0.001) (Fig. 3D). Immunohistochemistry showed expression of FABP6 and MTTP, two lipid transport proteins, in the upper glands of Mta2^cKO^ colon (Fig. 3G, 3H). Alkaline phosphatase, a small intestine brush border enzyme, was also activated and localized to surface colonocytes (Fig. 3G). Consistent with these molecular changes, BODIPY stain showed lipid accumulation in ileal villi and the upper glands of Mta2^cKO^ proximal colon, but not in control colon (Fig. 3I). Thus, MTA2 loss in colonocytes activated many small intestinal genes, particularly those involved in lipid transport and metabolism.

**Figure 3.**
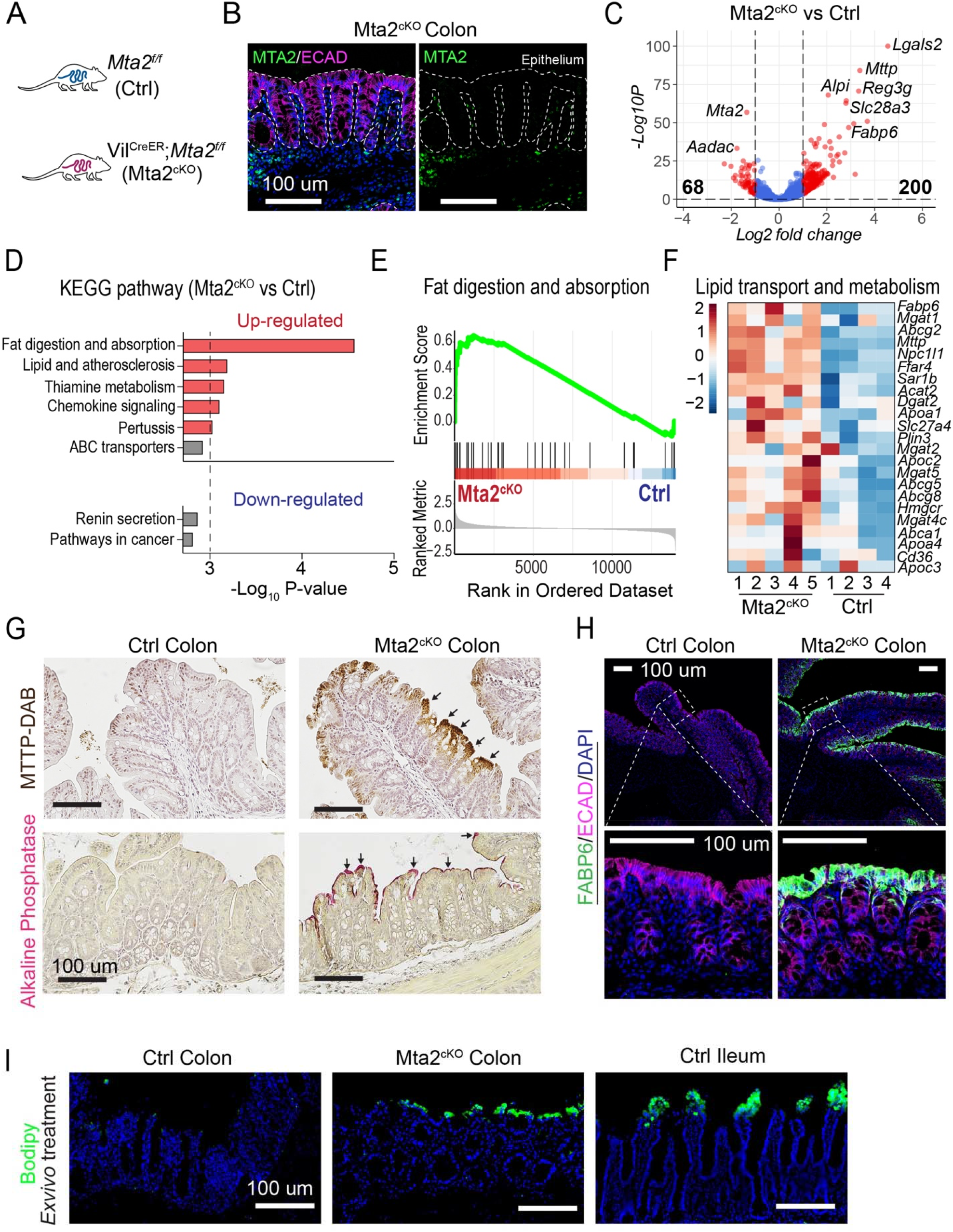
Activation of lipid transport and metabolism genes in adult mouse colon after MTA2 loss. (**A, B**) *Mta2* was deleted from 2-month old *Vil*^*CreER*^*;Mta2*^*f/f*^ (Mta2^cKO^) mice by applying tamoxifen (**A**), leading to near complete absence of MTA2 in colonic epithelium (**B**). (**C-F**) RNA-seq of control and Mta2^cKO^ colonic glands identified 200 up-regulated and 68 down-regulated genes ([LFC] > 1, adjusted p [padj] < 0.05) (**C**, volcano plot). Although no molecular pathways were significantly enriched among the down-regulated cohort, genes involved in lipid absorption, transport, and metabolism were prominently enriched among the up-regulated cohort, illustrated by KEGG pathway analysis (**D**) and GSEA (**E**) and in heatmap representation (**F**). (**G, H**) Histology and immunofluorescence staining showed activation of lipid transport proteins FABP6 and MTTP and small intestine brush border enzyme Alkaline Phosphatase in the surface colonocytes of Mta2^cKO^ colon. (**I**) BODIPY stain revealed presence of lipid accumulation in villi of ileum and surface glands of Mta2^cKO^ proximal colon, but not control colon.

### MTA2 retains HNF4A on colonic enhancers and prevents HNF4A from activating small intestine chromatin

MTA2 is part of the NuRD complex, which has been proposed to suppress alternative transcriptional programs in several tissues by direct binding and suppression of target genes^27^. To investigate how MTA2 modulates small intestine gene expression in colon, we mapped genome-wide MTA2 binding by chromatin immunoprecipitation sequencing (ChIP-seq). Duplicate MTA2 ChIP data from colonic epithelia yielded highly concordant data with 23,557 peaks (q < 1 ×10^−3^, using input DNA and Mta2^cKO^ as controls) (Fig. S4A). Colonic MTA2 binding occurred at promoters (49.1%, < 2kb from transcription start sites (TSSs)) and distal elements (50.9%, introns and intergenic regions) (Fig. S4B). Genes near MTA2 binding sites (< 50kb) were highly enriched for the colonic but not the small intestine signature (Fig. 4A). For instance, small intestine genes such as *Lgals2, Npc1l1, Abcg8*, and *Pla2g2a*, which are activated in *Mta2* null colon (Fig. 3F), lacked nearby MTA2 binding. These data indicate that MTA2 does not directly bind and regulate small intestine genes.

**Figure 4.**
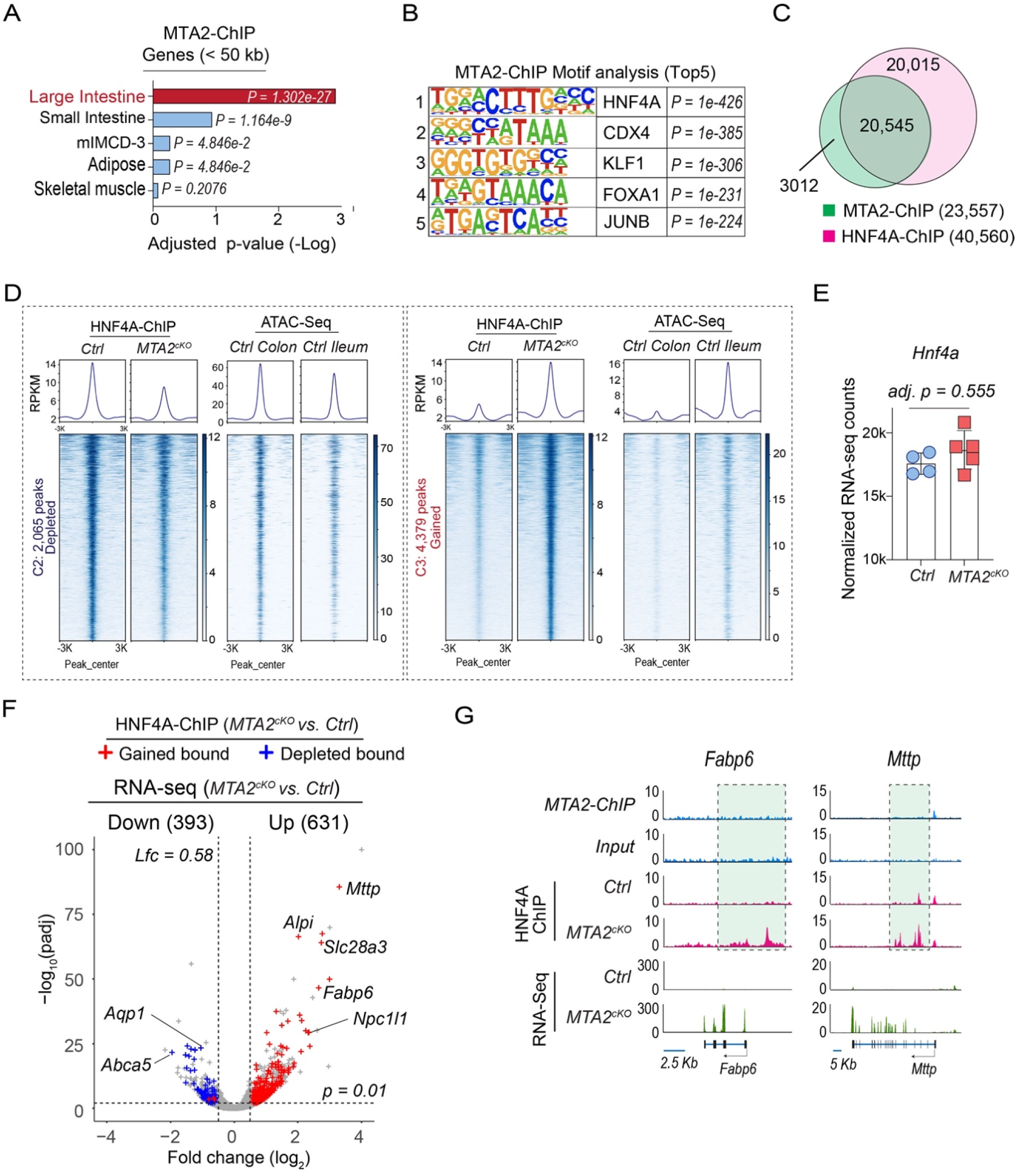
MTA2 retains HNF4A on colonic enhancers and prevents HNF4A from activating small intestine chromatin. **(A)** Tissue enrichment scores of genes near MTA2 binding peaks (MACS2, q <0.001, distance < 50 kb) in colon showed a predominant colonic signature. **(B)** Top 5 DNA binding motifs in MTA2 distal binding sites ranked by P value. **(C)** Venn diagram showing the overlap of MTA2 and HNF4A genomic binding in colon. **(D)** DNA binding profiles of HNF4A sites that were either reduced (left panel, 2,065 sites) or gained (right panel, 4,379 sites) after MTA2 loss in colon. Corresponding ATAC signals in colon or ileum were shown. Plots are 6-kb windows centered on each HNF4A binding site. **(E)** HNF4A mRNA levels were comparable in Mta2^cKO^ vs control colon. Mean ± S.D. N = 4-5 mice. Adjusted p value by Wald test corrected for multiple testing with Benjamini and Hochberg method. **(F)** Combined RNA-seq and HNF4A ChIP plot show that after MTA2 deletion, the up or down of gene expression is strongly associated with gain or loss of HNF4A binding. **(G)** Genome Browser tracks of MTA2 and HNF4A binding and RNA-seq at genomic loci of two small intestine lipid transport genes. Gain of HNF4A binding at these loci (highlighted) correlated with transcriptional activation.

Using HOMER analysis, we identified the DNA-binding motif of the intestinal transcription factor HNF4A as the top enriched motif at distal MTA2 binding sites (Fig. 4B). HNF4A and its homolog HNF4G are expressed in both small and large intestines and shown to be important in activating enterocyte gene transcription^28^, prompting us to evaluate whether MTA2 could regulate HNF4A in colon. Indeed, 87.2% of the MTA2 binding sites on colonic chromatin overlapped with binding of HNF4A (Fig. 4C). HNF4A expression was unchanged after MTA2 loss but HNF4A binding was depleted at 2,065 and acquired at an additional 4,379 sites in Mta2^cKO^ colon (log2FC > 1.0, q < 0.01) (Fig. 4D, 4E and S4D). About 80% of depleted sites (1,639 of 2,065) and nearly all gained sites (4,233 of 4,379) were in distal elements (Fig. S4E), indicating that MTA2 regulates HNF4A binding primarily at distal enhancers. Consistent with this notion, the depleted and gained HNF4A sites corresponded to areas of open chromatin enriched in colon and ileum respectively (Fig. 4D). Moreover, loss and gain of HNF4A binding in Mta2^cKO^ colon were strongly associated with down-regulation of colonic and up-regulation of ileal genes, respectively (Fig. 4F, 4G and S4F, S4G). Thus, MTA2 deletion led to HNF4A loss on colonic enhancers and its relocation to small intestine enhancers, triggering activation of small intestine genes in the colon. These data suggest that MTA2 retains HNF4A binding on colonic chromatin and prevents HNF4A from activating small intestine genes.

### Both SATB2 and MTA2 co-localize with HNF4A on colonic chromatin but SATB2 restrains HNF4A more strongly than MTA2

Both SATB2 and MTA2 can regulate colonic plasticity and our findings indicate that they interact physically (Fig. 1A-C). Structural studies of SATB1/2 proteins have identified four functional domains: a N-terminal ubiquitin-like domain (ULD) that mediates oligomerization, a CUT-like domain (CUTL) and two CUT domains (CUT1 and CUT2) that are critical for DNA binding, and a c-terminal HOX domain^29^ (Fig. 5A). Curiously, although HOX domains often serve as a primary DNA binding domain, this is not the case for SATB1/2^30^. The primary function of the HOX domain in SATB1/2 is unclear. We generated five SATB2 mutant proteins, each lacking one of the five functional domains (Fig. 5A). Co-IP studies with these mutant SATB2 proteins revealed that MTA2 interacts with SATB2 primarily via the HOX domain (Fig. 5B).

**Figure 5.**
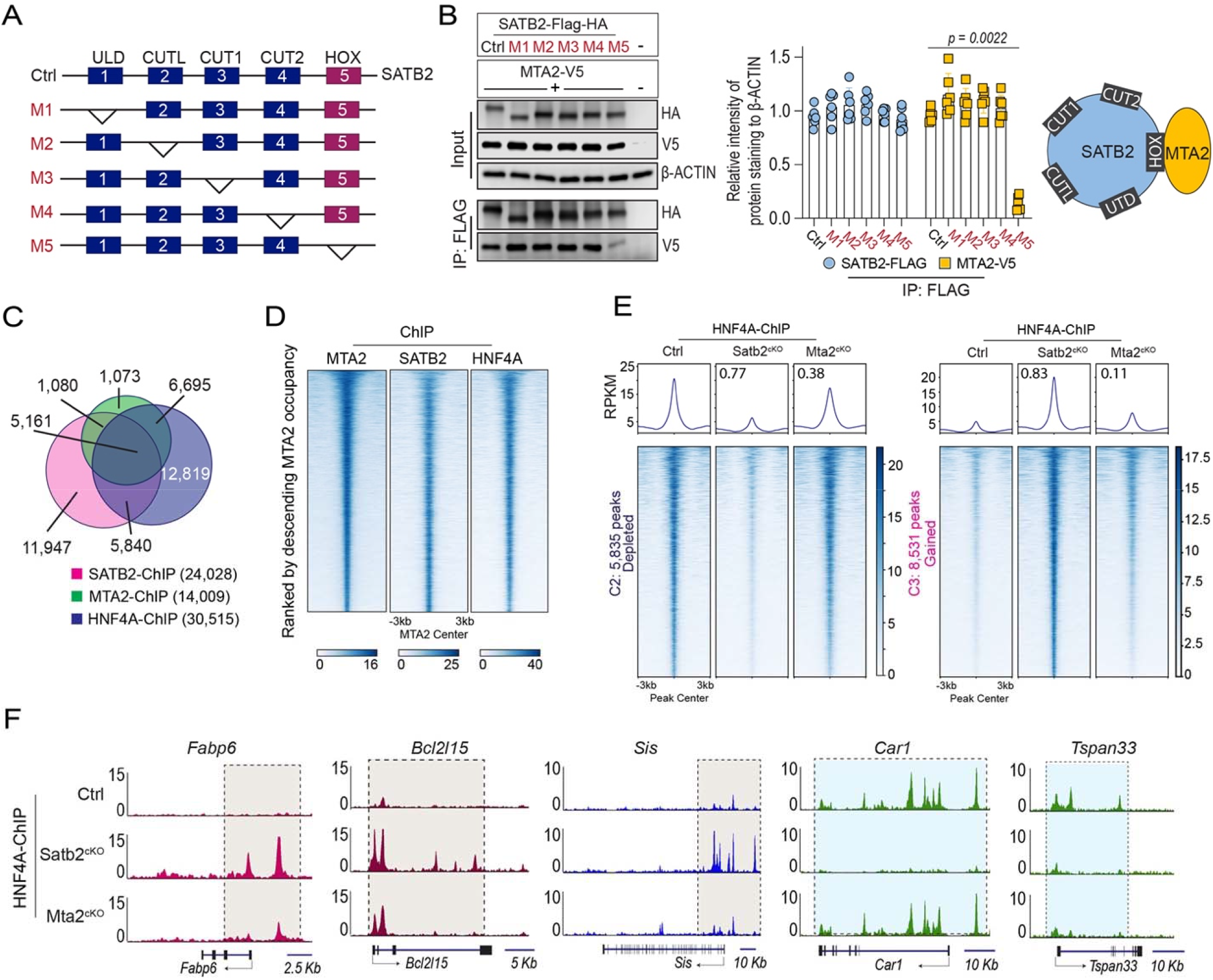
SATB2 and MTA2 co-bind with HNF4A on colonic chromatin but SATB2 retains HNF4A more strongly than MTA2. (**A, B**) We generated 5 mutant SATB2 proteins (M1-5) with each lacking one of the 5 functional domains (**A**). Co-IP of the SATB2 mutants and MTA2 showed that the SATB2 HOX domain was required for SATB2 interaction with MTA2; without the HOX domain, the interaction was abrogated (**B**). Mean ± S.D. n = 6. P value by unpaired t-test. (**C, D**) Overlap of SATB2, MTA2 and HNF4A distal genomic binding sites in colonic tissues was shown in the Venn diagram (**C**) and the DNA binding profiles (**D**). Peaks were ranked by descending MTA2 occupancy. **(E)** DNA binding profiles of HNF4A sites that were either reduced (left panel, 5,835 sites) or gained (right panel, 8,531 sites) in Satb2^cKO^ colon. In comparison, HNF4A loss or gain at these sites were modest in Mta2^cKO^ colon, but nonetheless statistically significant by Kolmogorov-Smirnov test (K-S D values shown in the density plots). Peaks centered on HNF4A binding sites in 6kb windows. **(F)** Genome Browser tracks of HNF4A binding at genomic loci of the small intestine genes *Fabp6, Bcl2l15* and *Sis* and the colonic gene *Car1* and *Tspan33* in Mta2^cKO^ and Satb2^cKO^ colon. SATB2 can more strongly influence HNF4A binding than MTA2.

The physical interaction of MTA2 and SATB2 suggests that both might co-localize with HNF4A on colonic chromatin. Because both MTA2 and SATB2 regulate HNF4A primarily at distal genomic sites (this study and reference 21), we assessed distal co-localization of the three factors. Alignment of MTA2 peaks with published SATB2 ChIP data showed 44.5% co-occupancy at distal elements, whereas 36.8% of distal MTA2 sites were co-bound by both SATB2 and HNF4A (Fig. 5C, 5D). Despite this extensive co-localization, SATB2 loss in colon activated more small intestine genes than MTA2 loss and larger numbers of HNF4A binding sites were lost and gained in Satb2^cKO^ (*Villin-Cre*^*ER*^; *Satb2*^*fl/fl*^) than in Mta2^cKO^ colon, with lost and gained sites corresponding to sites of open chromatin in colon and ileum respectively (Fig.5E, 5F and S5A). MTA2 genomic binding changes paralleled those of HNF4A in *Satb2* null colon (Fig. S5A, S5B). Altogether, these data suggest that both MTA2 and SATB2 could restrain HNF4A at colonic chromatin, but SATB2 retains HNF4A more robustly.

### Small intestine gene activation in both Mta2^cKO^ and Satb2^cKO^ colon depends on HNF4A

Our chromatin mapping data implicated HNF4A as a key mediator of colon-small intestine plasticity and gain of HNF4A binding on small intestine chromatin is tightly associated with transcriptional activation of small intestine genes in Mta2^cKO^ or Satb2^cKO^ colon (this study and reference 21). If this association is causal, then removing HNF4A should block colonic transcriptional plasticity. To evaluate this hypothesis, we differentiated cultured mouse colonic organoids into colonoids enriched for CA1^+^ colonocytes (Fig. 6A and 6B). We used CRISPR-Cas9 to delete *Hnf4a* from either Mta2^cKO^ or Satb2^cKO^ colonoids, achieving deletion efficiencies > 90% (Fig. 6C and 6E). qPCR analysis of small intestine genes showed that their activation was attenuated in both mutants (Fig. 6D and 6F).

**Figure 6.**
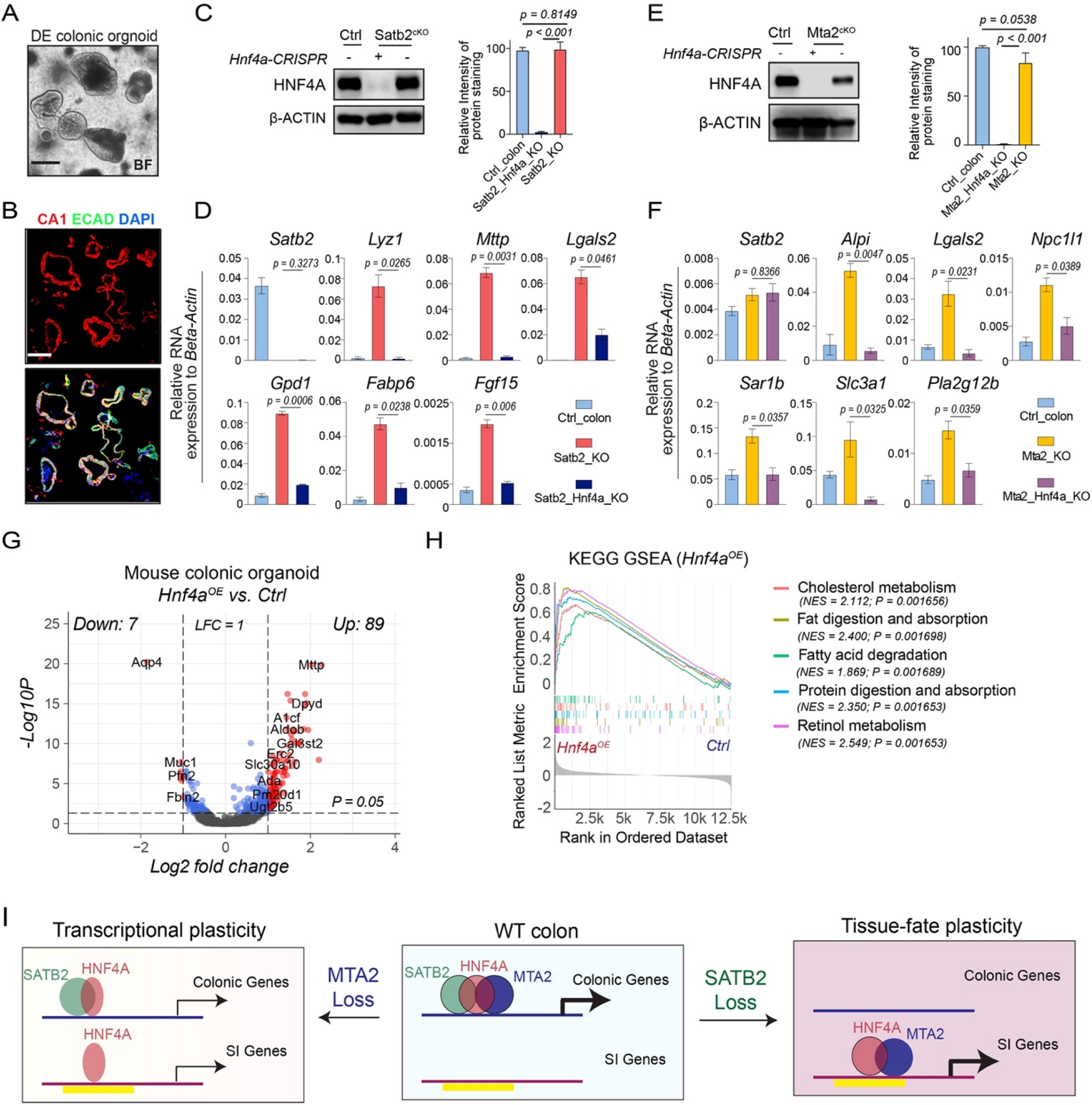
Activation of small intestine genes in both Mta2^cKO^ and Satb2^cKO^ colon is dependent on HNF4A. (**A, B**) Bright field and immunofluorescence pictures of differentiated (DE) colonic organoids. CA1 staining showed enrichment of colonocytes in these organoids. BF: bright field. **(E)** Immunoblot and quantification showed that HNF4A levels were comparable in wild-type (WT) vs Satb2^cKO^ colonoids. CRISPR reduced HNF4A in Satb2/Hnf4a double knockout colonoids to less than 5% of control levels. Mean ± S.D. n = 3. P value by Unpaired t-test. **(F)** QPCR showed that loss of HNF4A blocked small intestine gene activation in *Satb2* mutant colonoids. **(G)** Immunoblot and quantification showed that HNF4A levels were comparable in wild-type (WT) vs Mta2^cKO^ colonoids. CRISPR reduced HNF4A in Mta2^cKO^ colonoids to less than 2% of control levels. n = 3. P value by Unpaired t-test. **(H)** Loss of HNF4A attenuated small intestine gene activation in *Mta2* mutant colonoids. *Satb2* mRNA level was not altered by *Hnf4a* deletion. (**G, H**) Over expression of HNF4A (Hnf4a^OE^) in colonic organoids led to mostly up-regulation of small intestine genes (**G**) enriched for pathways characteristic of small intestine functions (**H**). (**I**) A proposed model of colonocyte plasticity regulation in which MTA2 and SATB2 form a chromatin complex at colonic chromatin to retain HNF4A. SATB2 restrains HNF4A more tightly than MTA2. MTA2 loss leads to a modest depletion of HNF4A on colonic and gain on small intestine (SI) chromatin, and modest down- and up-regulation of colonic and small intestine (SI) genes. In contrast, SATB2 loss results in the untethering of large numbers of HNF4A and consequently transcriptomic shift from colon to small intestine. Yellow bar denotes primed small intestinal enhancers in colon.

Given that colonic enhancers are occupied by HNF4A whereas the small intestine enhancers are primed but lack HNF4A binding in colon, we reasoned that ectopically expressed HNF4A might engage small intestine enhancers in colon and activate transcription. To test this hypothesis, we over-expressed HNF4A in cultured colonic organoids. RNA-seq showed 89 up-regulated and 7 down-regulated genes (LFC > 1, p < 0.05) (Fig. 6G and Fig. S6). Gene sets characteristic of small intestine functions such as cholesterol metabolism, fat and protein digestion and absorption, and retinol metabolism, were enriched among the up-regulated genes (Fig. 6H). Thus, both loss- and gain-of-function studies support HNF4A as a central mediator of small intestine gene activation in colonic plasticity. Collectively, our data support a model of colonic plasticity regulation in which the chromatin factors MTA2 and SATB2 form a complex at colonic chromatin to retain HNF4A. The degree of plasticity depends on the amount of HNF4A released from this restraint. If relatively little HNF4A is liberated as with MTA2 loss, then a modest increase of small intestine genes results; if large numbers of HNF4A are released as with the SATB2 loss, then transcriptomic shift and overt phenotypic conversion ensues (Fig. 6I).

## Discussion

To perform specialized functions, distinct cell types must maintain unique identities, including cell type-specific transcriptomes. They also need to adapt to changing environments by deploying plasticity in transcription or even cell identity^31-33^. The molecular control of cell identity and plasticity is a fundamental question, with scant mechanistic insights.

We previously uncovered surprising plasticity between adult colon and ileum, controlled by the colon-restricted chromatin factor SATB2. SATB2 deletion causes drastic cell fate switch, converting colonocytes to enterocytes. In this study, we identify MTA2 as a new component of the molecular machinery that preserves colonic identity and mediates plasticity. MTA2 loss led to activation of a subset of ileal genes and modest down-regulation of colonic genes. Thus, MTA2 and SATB2 regulate colonic plasticity in different degrees. Nevertheless, a common underlying mechanism is the ability of MTA2 and SATB2 to retain HNF4A on colonic chromatin. SATB2 appears able to “tether” HNF4A strongly to colonic enhancers, hence its loss leads to transcriptome-wide changes, whereas MTA2 tethers a fraction of HNF4A and its loss causes more modest transcriptional change. Notably, small intestine gene activation in both MTA2 and SATB2 knockout colon is blocked by removing HNF4A, highlighting the central role of this transcription factor in mediating colonic plasticity.

MTA2 is part of the NuRD complex, which alters nucleosome spacing and deacetylates histones and other proteins^34-37^. Incapacitating NuRD in multiple tissues, including lymphoid cells and muscles, activates alternative lineage programs^38,39^. However, in these cases, NuRD is proposed to directly bind and silence alternative lineage genes, a mechanism distinct from the one we observed in colonic to ileal plasticity. The exact mechanism by which MTA2 and SATB2 retain HNF4A on colonic chromatin has yet to be fully understood. We speculate that loss of MTA2 changes the local chromatin milieu, making it less favorable for HNF4A binding. This notion is consistent with studies in embryonic stem cells where NuRD binding on active enhancers is reported to reduce or enhance TF binding depending on the context^40^. As SATB2 likely engages multiple chromatin remodeling factors in addition to MTA2, its loss shifts the colonic transcriptome more profoundly. In sum, our study has begun to reveal a chromatin complex that serves the dual purpose of maintaining colonic identity and mediating plasticity. We propose that this model of cellular plasticity may apply broadly, that is, chromatin complexes restrain critical TFs at tissue-specific chromatin to prevent them from activating alternative lineages; measured release of these factors leads to different degrees of cellular plasticity.

---

**Supplementary Figure 1.**
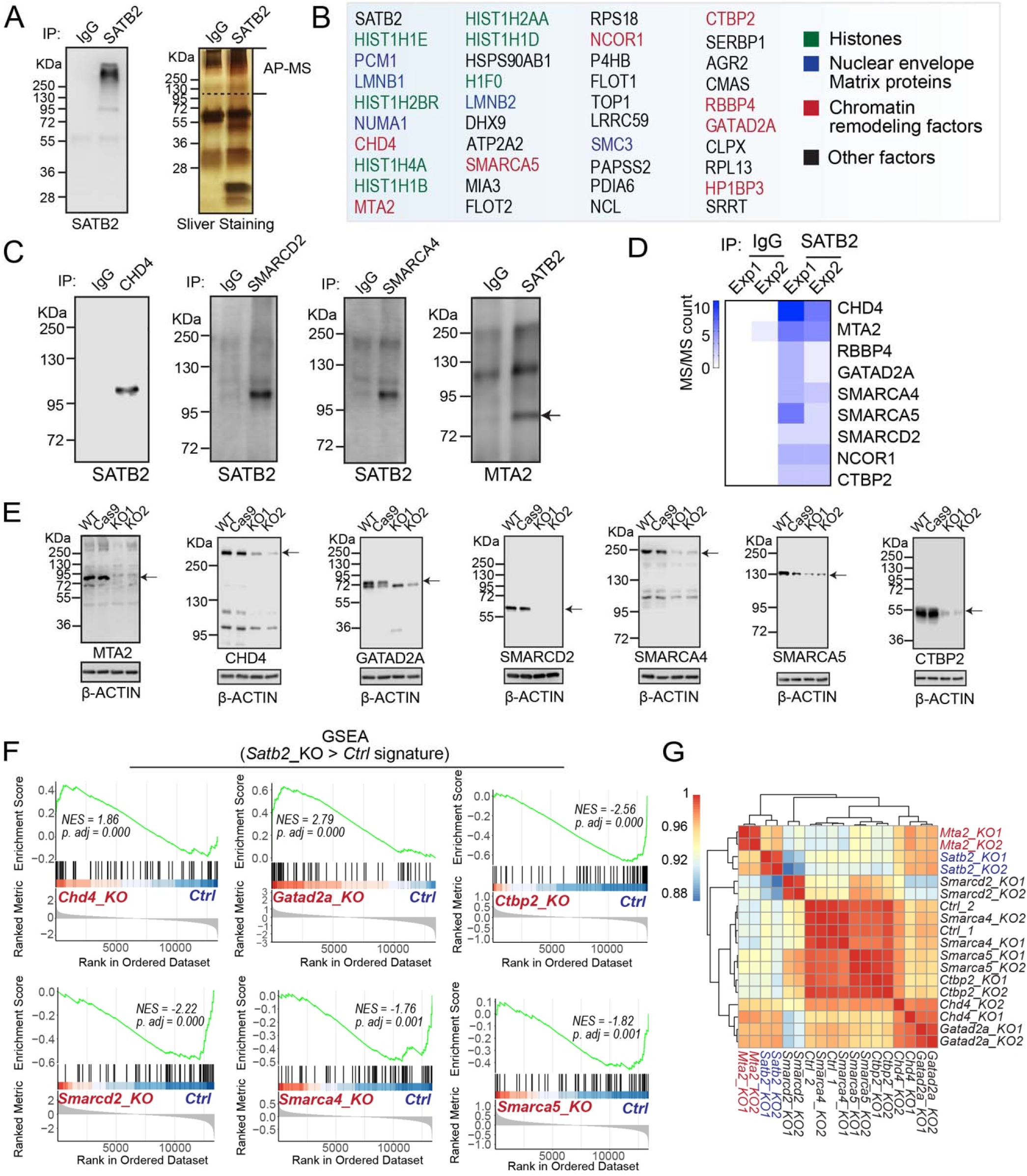
AP-MS and CRISPR screen identified candidate SATB2-associated factors that regulate colonic transcription. **(A)** Immunoblot of cross-linked murine colonic tissues probed with anti-SATB2 antibody revealed protein complexes at high molecular weight (left panel). Right panel shows silver stain of protein gel loaded with control IgG or anti-SATB2 immunoprecipitated colonic protein samples. The high molecular weight portion of the gel (above the dotted line) was excised for Mass Spectrometry. **(B)** The top 40 enriched proteins from two independent AP-MS experiments. **(C)** Co-IP with murine colonic tissues showed physical interactions between SATB2 and several of the chromatin remodeling factors identified in AP-MS. **(D)** A heatmap of MS counts of the nine chromatin remodeling factors that we selected for CRISPR study. **(E)** We used CRISPR-CAS9 to disrupt seven chromatin remodeling factors in cultured murine colonic organoids. Immunoblot of two independent experiments, together with two different controls (unmanipulated colonic organoids or colonic organoids transduced with CAS9 but not gRNA), showed successful disruption of each factor. **(F)** GSEA analysis of RNA-seq data from colonic organoids in which different chromatin remodeling factors were knocked out. Genes up-regulated in Satb2 knockout vs control colonic organoids was used as the gene set for comparison. Positive correlation was seen for Chd4 and Gatad2a, but not Smarcd2, Smarca4, Smarca5, or ctbp2, knockout organoids. **(G)** Pearson correlation plot of transcriptomes from different CRISPR knockout organoids and controls. Mta2 knockout transcriptome had a stronger correlation with Satb2 knockout transcriptomes than the other samples.

**Supplementary Figure 2.**
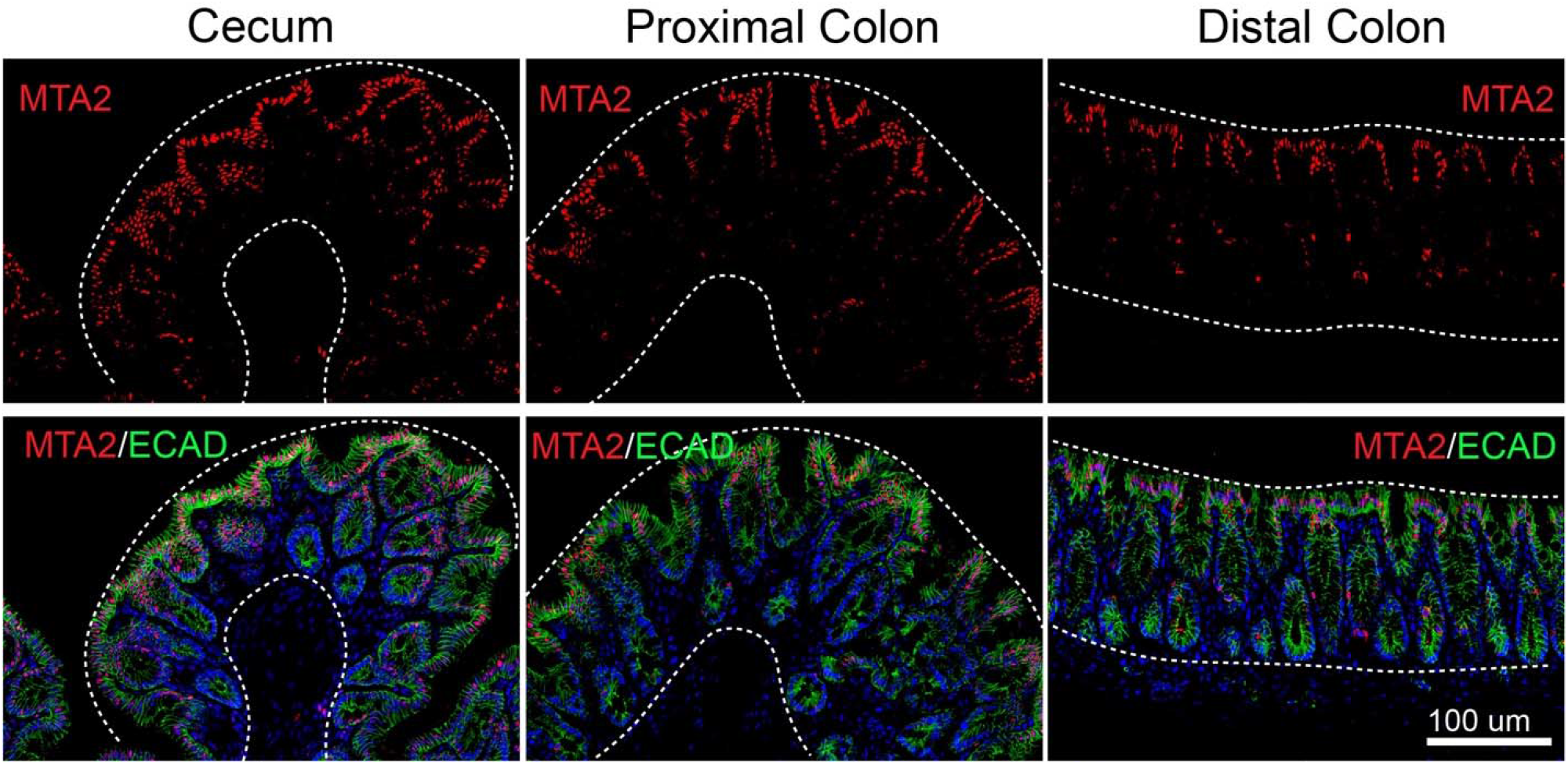
MTA2 is expressed in the upper glands of adult mouse colon. Immunofluorescent staining of MTA2 in different regions of adult mouse large intestine showed a predominant localization in the upper glands. Some non-epithelial cells also expressed MTA2.

**Supplementary Figure 3.**
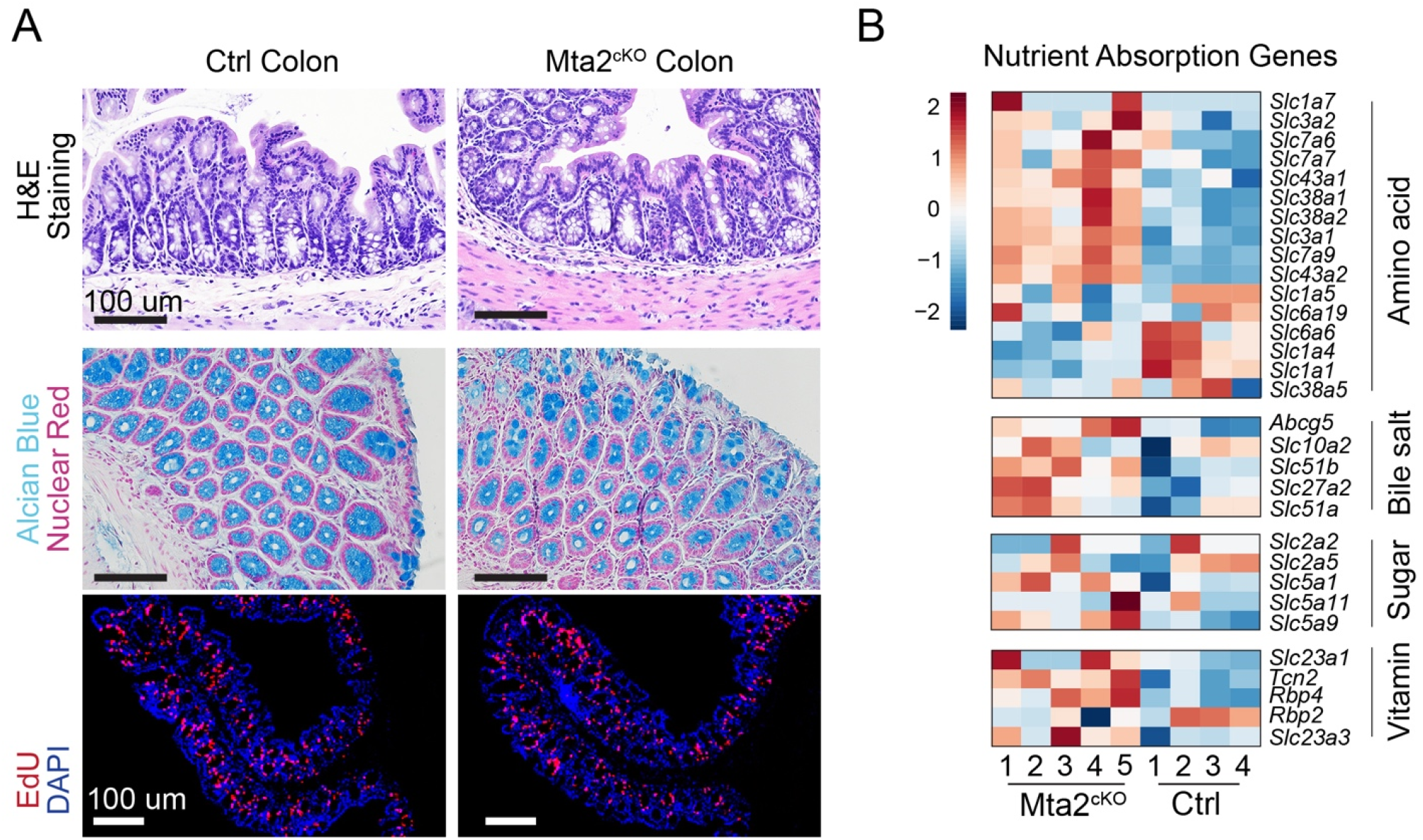
Analysis of Mta2^cKO^ colon. **(A)** Mta2^cKO^ colon appeared normal compared with controls, based on H & E stain, Alcian blue stain for goblet cells, and EdU labeling of the proliferative zone. **(B)** Heat map of normalized RNA-seq counts of nutrient transporter genes in control and Mta2^cKO^ colon.

**Supplementary Figure 4.**
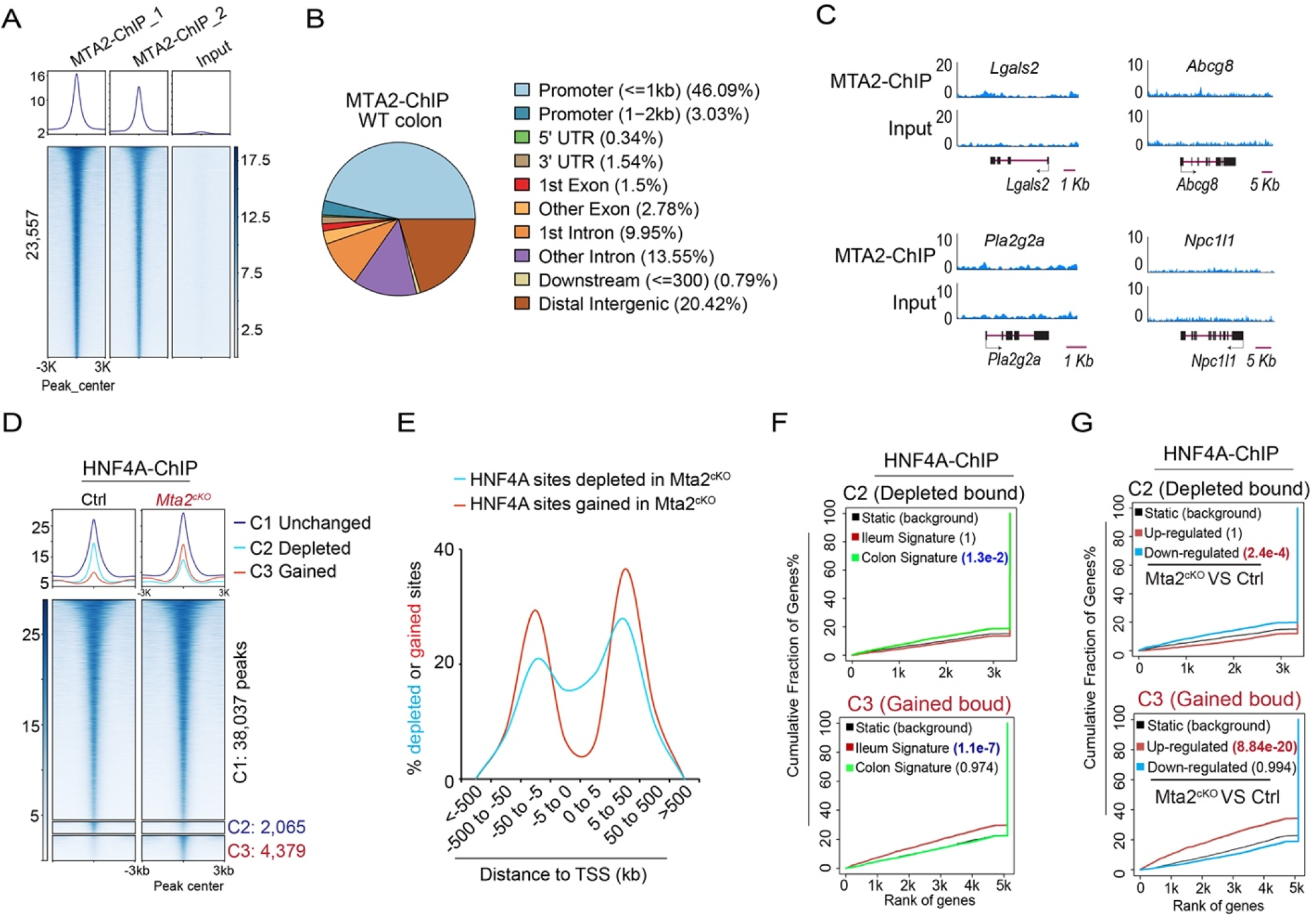
MTA2 regulates HNF4A binding in colon. (**A, B**) MTA2 genomic binding sites in colonic tissues were identified by ChIP-seq using input DNA as controls, yielding 23,557 peaks (peak call by MACS2, duplicate biological samples) (**A**). The binding sites include distal elements as well as promoters (**B**). **(C)** Genome Browser tracks showing a lack of MTA2 binding at genomic loci of 4 small intestine genes. **(D)** DNA binding profiles of HNF4A sites that were unchanged (C1, 38,037 sites), reduced (C2, 2,065 sites), or gained (C3, 4,379 sites) after MTA2 loss in colon. Plots are 6-kb windows centered on each HNF4A binding site. **(E)** DNA binding density plots of HNF4A sites that were depleted (2,065 sites) or gained (4,379 sites) after MTA2 loss in colon. TSS: translation start site. (**F, G**) Predictions of enhancer regulatory functions by BETA (binding and expression target analysis) indicate that loss of HNF4A binding in Mta2^cKO^ colon is associated with down-regulation of colonic genes (**E, F**, upper plots) whereas gain of HNF4A binding is associated with up-regulation of ileal genes (**E, F**, lower plots). Plots depicts the cumulative score of regulatory potential for every gene based on enhancer distances from the TSS. Blue lines represent the background of unaltered genes, and p values denote the significance of up or down associations relative to the background.

**Supplementary Figure 5.**
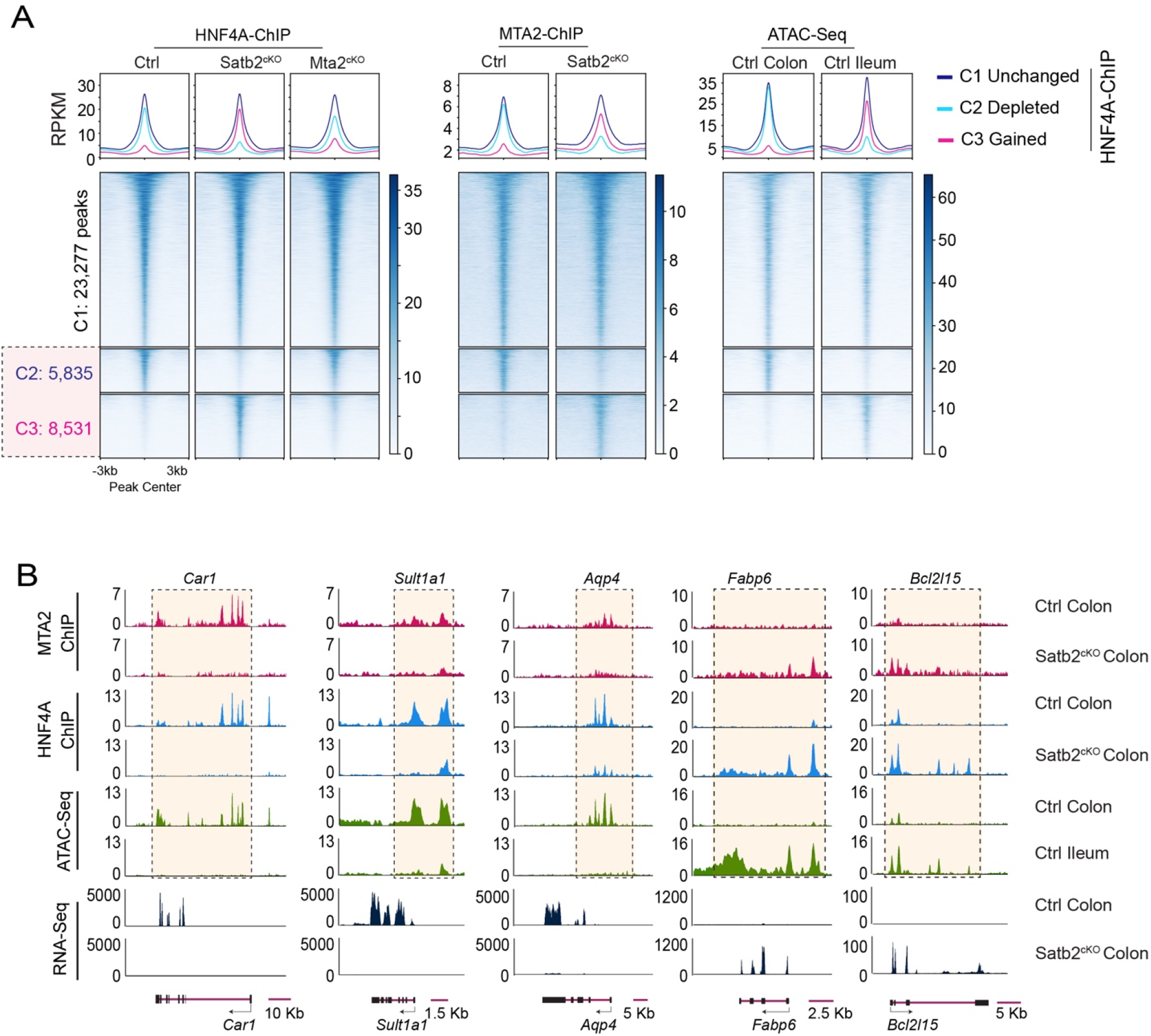
Differential regulation of HNF4A binding by SATB2 vs MTA2. **(A)** HNF4A DNA binding profile comparison in Satb2^cKO^ vs Mta2^cKO^ colon. Lost and gained sites (C2 and C3 respectively) were highlighted. Corresponding MTA2 binding profiles and ATAC profiles in wild-type colon and ileum were shown. Larger HNF4A binding changes were seen in Satb2^cKO^ than Mta2^cKO^ colon. MTA2 binding changes parallel that of HNF4A in Satb2^cKO^. Plots are 6-kb windows centered on each MTA2 binding site. **(B)** Genome Browser tracks showed coordinated shifts in MTA2 and HNF4A binding, ATAC and mRNA profiles at genomic loci of three colonic genes (Car1, Sult1a1, and Aqp4) and two small intestine genes (Fabp6 and Bcl2l15) before and after *Satb2* deletion.

**Supplementary Figure 6.**
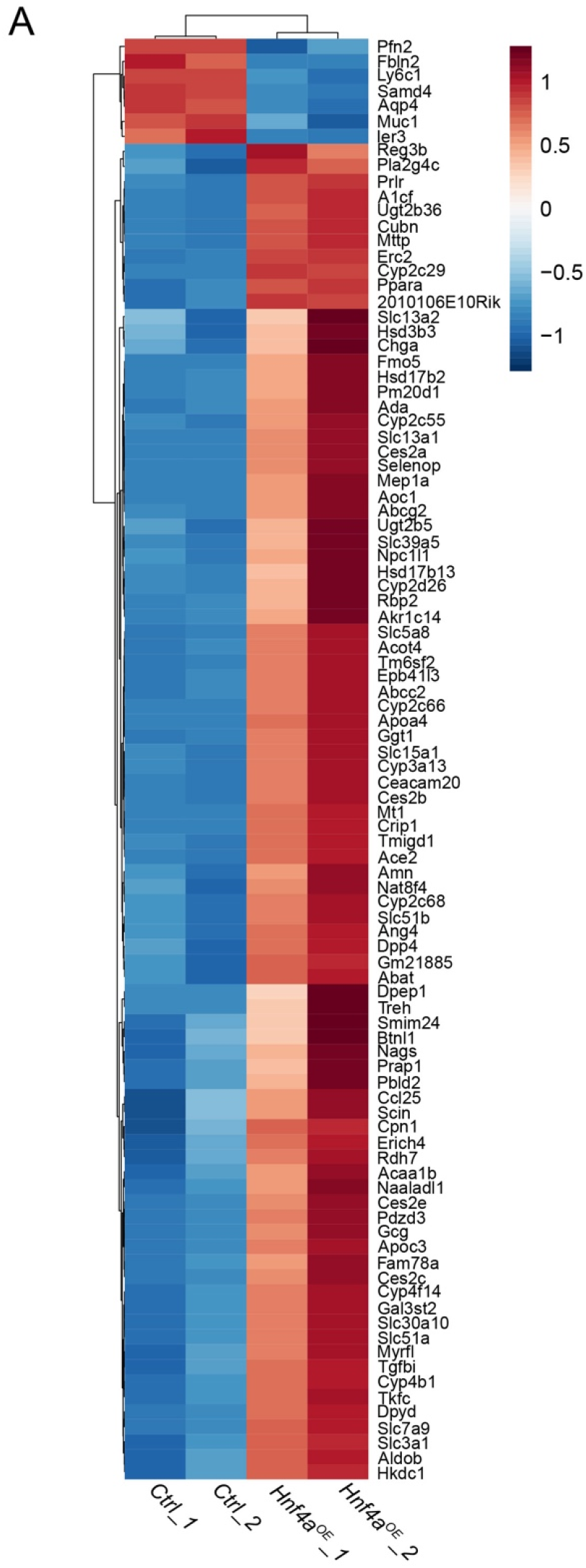
Ectopic HNF4A expression in mouse colonic organoids activated small intestine genes. Heatmap of all differentially expressed genes (LFC > 1, P < 0.05) in Hnf4a over-expression (Hnf4a^OE^) colonic organoids by RNA-seq.

## Acknowledgements

We thank the WCM Epigenetics Core (Jenny Xiang) for excellent technical assistance. We are grateful to Sean Houghton and Shariq Madha for assistance with computational analyses, Drs. Robert Roeder and Yi Zhang for providing Mta2^f/f^ mice. This work was supported by a Tri-Institutional Stem Cell grant to Q.Z and a NIH/NIDDK grant to Q.Z and R.S. (1R01DK125817).

## Author contributions

W.G and Q.Z devised the experiments, interpreted data and wrote the manuscript. W.G performed majority of the experiments with the help of X.H, Y.L and J. M. G. X.H analyzed RNA-seq data. P.N.P.S analyzed ChIP-seq data. S.R and M.V provided advice and data. R.S advised the project and edited manuscript. Q.Z. supervised the project.

## Methods

### Resource Availabilit2

#### Lead contact

Further information and requests for resources should be directed to the Lead Contact, Qiao Zhou

#### Materials Availability

All the materials will be available upon request to corresponding author under material transfer agreement with Weill Cornell Medicine.

#### Data and Code Availability

The High-through sequencing raw and processed data have been deposited to Gene Expression Omnibus (GEO). Bulk RNA-Seq: GSE213879 and GSE213878. ChIP-Seq: GSE213877. We also analyzed our previously published GEO datasets: GSE148690, GSE167283, GSE167287, and GSE167284 (Gu et al.,2022).

### Mouse strains

All mouse experiments were conducted under the IACUC protocol 2018-0050 at Weill Cornell Medical College. The MTA2loxp/loxp (MTA2f/f) strain (Xiangdong Lu et al. 2008) was a gift from Dr. Robert G. Roeder of the Rockefeller University and Dr. Yi Zhang of Harvard Medical School who originally made the strain. Vil-CreERT2 strain (el Marjou et al., 2004) was a gift from Sylvie Robine (Institute Pasteur). To confer conditional deletion of MTA2, 4 mg per 25 g of body weight of tamoxifen (TAM, 10 mg per ml in corn oil) was intraperitoneally injected into both Mta2^cKO^ (Vil-CreERT2; *Mta2*^f/f^) and Ctrl (*Mta2*^f/f^) mice (male and female equally, at 2 months old) once every 2 days for a total of 3 times. Mice were housed and breaded in a temperature- and humidity-controlled environment with 12 hours light/dark cycle and food/water supplement.

### Mouse colonic crypt isolation, colonoid culture and differentiation

Mouse colonoid line establishment and culture were performed as previously described with modification (Gu et.al, 2022). All the experiments were performed on ice or at 4°C unless specified. Briefly, proximal colon top glands were scraped by glass slides and then the tissues were cut into approximately 0.3 cm size pieces and incubated in 10 mM EDTA for 30 mins. The tissues were transferred to 15 mL cold PBS solution. After vigorous pipetting with 1% BSA pre-coated 10 mL serological pipettes, epithelium cell clumps were collected by centrifugation at 300g for 5 mins. Crypts were further isolated by filtering through a 70 μm cell strainer. 25-100 Crypts (P0) per 12 μl Matrigel™ droplet were cultured in WENR medium (Table S1) in humidified chambers containing 5% CO_2_ at 37°C for 4 -5 days with one-time medium change at day 2. After one passage, the P1 colonoids were differentiated into colonocyte enriched colonic organoids by culturing in differentiation medium (DEM) (WENR medium without WRN conditioned medium and with the addition of 1 μg/mL RSpondin and 10 μM L-161,982, Table S1) at day 2. Three days after differentiation, the organoids were either directly lysed in RNA lysis buffer (ZYMO) for RNA exaction, or incubated with cell recovery solution on ice, to remove Matrigel, for immunofluorescence, immunoblotting and immunoprecipitation analyses.

### CRISPR-mediated gene knockout in colonoids

Murine *Mta2, Chd4, Gatad2a, Smarca4, Smarca5, Smarcd2 and Ctbp2* sgRNAs were designed with the Synthego CRISPR design tool (Table S2) and cloned into a LentiCRISPRv2 vector (Addgene plasmid #52961). The lentiviruses were packaged with second-generation helper plasmids by transfection with lipofectamine 3000. Each virus titer was determined by counting puromycin resistant clones in HEK293T cells 5 days after infection.

To generate the colonoids with gene ablation, a total of 10^5^ cells suspensions (TypLE digested small colonic fragments with 1 to10 cells per fragment) in 200 μL WENR with 10 μg/mL polybrene were mixed with 20 μL of 10^8^ TCID_50_/ml of virus in one well of a non-tissue culture treated 24 well plate, and centrifuged at 1,100g at 37°C for 30 mins to facilitate infection. After centrifugation, 200 μL of WENR was added and the plate was further incubated for 4 hours at 37°C. Cells were then resuspended, pelleted, and embedded in Matrigel™ as described in coloniods culture method section. Puromycin selection (1 μg/mL) was initiated 3 days post infection and lasted for 4 days. After puromycin selection, colonic organoids were sub-cultured into new Matrigel drops and differentiated in DEM for 3 days. The CRISPR-mediated deletion efficiency was analyzed with immunoblotting by using specific target antibodies (Key Resource Table).

### Affinity Purification Mass Spectrometry (AP-MS)

All the experiments were performed on ice or at 4°C unless specified. Murine proximal colon tissues (half of colon length, about 40 mm) were flushed clean by cold PBS and cut into approximately 3 mm size pieces. The tissues were incubated with 10 mM EDTA/PBS in 50 mL tube for 30 mins and then transferred to 20 mL cold PBS. After vigorous pipetting with 1% BSA pre-coated 10 mL serological pipettes, the colon tissues were removed to a new tube containing 10 mM EDTA/PBS and incubated for another 30 mins. Epithelium glands in suspensions were collected by centrifugation at 300g for 5 mins as fraction one, followed by resuspending in 10 mL PBS containing 10 μM Y-27632. The second fraction of epithelium glands were collected as the same procedure for fraction one. Two fractions of EDTA stripped epithelium glands were combined, pelleted, and cross-linked with 2 mM disuccinimidyl glutarate (DSG, Thermo Fisher Scientific, 20593) in PBS at room temperature (RT) for 45 mins. Pellets of epithelial cells were incubated with 0.3M RIPA buffer (Table S3) supplied with Protease Inhibitor Cocktails and sonicated at 20% amplification for 1 min (20 sec on and 20 sec off, 3 cycles). After 10 mins of centrifugation at 18,000 g, supernatants were collected and incubated with anti-SATB2 (Key Resource Table) overnight with a rotation speed of 10 RPM. After adding 30 μl protein A/G magnetic beads for 90 mins on the next day, the protein and beads complex was pulled down by a magnetic stander. Total 6 wash of 0.3M RIPA buffer were performed. Then the DSG-crosslinked SATB2 interaction protein complexes were cleaved from Protein A/G beads by boiling in Laemmli Sample Buffer from Bio-Rad without adding reducing reagents (DTT or 2-Mercaptoethanol) and separated by SDS-PAGE. Proteins in gel was visualized with sliver staining kit from Sigma. The gels over 100 KDa (SATB2 molecular weight) for each sample were cut and digested with trypsin, followed by desalting and LC-MS/MS for protein identification and quantification. The data were processed by MaxQuant and searched against Uniport mouse database with a 1% false discovery rate. Significantly enriched genes were filtered by the quadrilaterals that both samples have SATB2 AP-MS signal intensity and MS count over IgG controls >= 2 fold changes.

### Dual Cross-linking ChIP-Seq

ChIP for Transcription Factors (TFs) SATB2, HNF4A, and MTA2, was performed as described (Saxena et al., 2017; Gu et al., 2022). EDTA stripped primary colon glands from Ctrl, MTA2^CKO^ and SATB2^CKO^ were pelleted and cross-linked with 2 mM DSG at RT for 45 mins, followed by 1% formaldehyde fixation for 10 mins. For each experiment, about 50 μl of pelleted cross-linked cells were resuspended in 350 μl sarkosyl lysis buffer (0.25% sarkosyl, 1 mM DTT and Protease Inhibitor Cocktails in 0.3M RIPA buffer and sonicated at 60% amplification by a tip sonicator (125W) for 6 mins (45 sec on and 45 sec off, 8 cycles) to obtain 200bp to 800bp chromatin fragments. Lysates were spun down at 20,000g at 4°C to remove insoluble fractions. The supernatant was further diluted in 0.3M RIPA buffer with Protease Inhibitor Cocktails in a final 2 ml volume. Diluted lysates were incubated with TFs antibodies (Key Resource Table) at 4°C overnight and were additionally incubated with 30 μl protein A/G magnetic beads for 90 mins the next day. A total of 6 washes with cold 0.3 M RIPA buffer and final rinse with TE buffer (PH 7.5) were performed to remove nonspecific binding. Dual cross-links were reversed overnight by incubating at 65°C in 1% SDS and 0.1 M NaHCO_3_. Any remaining proteins were digested by Proteinase K for 1 hour at 37°C. Pulled down genomic DNA was purified with a MinElute purification kit and quantitated by Qubit. The libraries were prepared using the ThruPLEX DNA-Seq Kit, followed by a size selection and purification (200bp to 800bp, including index) with AMPure XP beads. The final libraries were quality controlled and pooled for sequencing.

### ChIP-Seq analyses

ChIP-seq libraries (MTA2, HNF4A and SATB2) were sequenced on an Illumina 4000 instrument to obtain 50-bp pair-end reads. All reads were trimmed (https://www.bioinformatics.babraham.ac.uk/projects/trim_galore/), and subject to quality control with FastQC before and after adapter trimming. ChIP-Seq analyses, including alignments, bigwig generation, reads mapping, peak calling, quantitative comparisons, peak annotation, and enriched motifs analysis were performed as described previously (Gu et al., 2022). BETA (Wang et al. 2013) was used to associate genes with HNF4A-ChIP depleted or gained bound sites and quantify these associations using peaks within 50-kb from TSS, a significance threshold of FDR-adjusted P < 0.01 for differential gene expression in wild-type ileum vs. colon or Mta2^cKO^ vs. Ctrl colon, and other default parameters.

### Edu labeling and Immunostaining

A Click-iT™ EDU Cell Proliferation Kit with Alexa Fluor® 555 (C10338) was used to evaluate proliferation. Briefly, 50 μg EDU per gram of mice bodyweight were intraperitoneal injected for 24 hrs. Mice were euthanized and then intestinal tissues were harvested and flushed clean with cold PBS. The tissues were cut into 1 cm pieces and fixed with 4% paraformaldehyde immediately at 4°C for 1 hour (Organiods were fixed for 20 mins). After wash with PBS, the tissues were dehydrated by 30% sucrose solution overnight and embedding in O.C.T for a Cryostat sectioning.

Immunofluorescence was performed using a standard procedure, incubating with primary antibodies (Key Resource Table) at 4°C overnight, followed with secondary antibodies at RT for 1 hr. The images were captured using either a confocal microscope (710 Meta) or a Nikon fluorescence microscope. For immunohistochemistry, samples were processed through heat mediated antigen retrieval in Citric Acid buffer (pH 6.0) for 15 mins. Samples were then stained with Anti-MTTP or MTA2 antibodies, followed by Goat anti-Rabbit HRP incubation at RT for 1 hr, and finally, developed with DAB (Brown color, Vector Laboratories, SK-4103) HRP Substrate. Alkaline phosphatase enzyme was detected by Stemgent AP staining kit 2 (Pink color). An Alcian Blue Stain Kit (Vector Laboratories, H-3501) was used to stain goblet cells.

### Western Blot

For Western blot analysis, monoclonal rabbit anti-SATB2 and SMARCD2, monoclonal mouse anti V5 and FLAG, polyclonal rabbit anti-CHD4, MTA2, CTBP2, SMARCA4, SMARCA5, GATAD2A, HDAC1 and HDAC2 antibodies were used to bind target protein, followed by an incubation with a secondary anti-Rabbit Peroxidase (HRP) or anti-Mouse HRP. Protein bands were visualized using enhanced chemiluminescent substrate (Pico from Thermo fisher) and recorded by a Li-COR C-Digit blot scanner. The relative signal intensity was quantified by Image J (v1.51 (100)).

### Bulk RNA sequencing analysis

Bulk RNA-seq was performed as previously described except for mapping to the mouse reference genome mm10 instead of mm9 (Banerjee et al., 2018). Briefly, reads alignment was performed by STAR package (Dobin et al., 2013). The raw count tables were generated by featureCounts (Liao et al., 2014). The DEseq2 package was used for differential expression analysis (Love et al., 2014). In DESeq2, the p-values attained by the Wald test are corrected for multiple testing using the Benjamini and Hochberg (BH) method. The Limma package (Ritchie et al., 2015) was used to remove donor-donor variance and batch-effect. Differentially expressed genes were generally determined using parameters of adjusted p-value < 0.05 and LFC > 1 or < -1 unless specified in figure legends. The heatmaps were plotted using the R package, pheatmap. GSEA KEGG analysis and GSEA analysis were conducted with the clusterProfiler package (Yu et al., 2012).

### BODIPY staining

Both Mta2^cKO^ and Ctrl mice were under high-fat diet treatment for 3 weeks. The mice were euthanized, about 2 cm size of colon segments were cut open and flushed with cold DPBS. Colon segments were then incubated with (10 μg/mL) BODIPY in DPBS at RT for 30 mins and followed with 3 times DPBS wash to remove excess BODIPY. Tissues were fixed with 4% paraformaldehyde immediately at 4°C for 1 hour. After wash with DPBS, the tissues were dehydrated by 30% sucrose solution at 4°C for 3 hours and embedding in O.C.T for a Cryostat sectioning.

### SATB2 domain deletion

pENTR-3xFlag-3xHA-mSATB2 and overlap PCR primers information were provided in Table S2. Each domain deletion PCR amplification was performed by TAKARA-HIFI Tag by following the manufactory guides. PCR fragments were then treated with Dpn1 at 37°C for 1 hour to remove plasmid template. After gel purified, the fragments were further phosphorylated by T4 PNK and self-ligated by Quick ligase. Each domain deletion cloning vector was confirmed by sequence analysis and recombined into a Dox-inducible Destination vector (Plx403) using Gateway LR Clonase II enzyme kit.

pENTR-mMTA2-V5 was cloned and recombined into Pinducer20. Plx403-SATB2 or each domain deletion expression vector was co-transfected with Pinducer20-MTA2 into 293T cells for 48 hours. Doxycycline (1 μg/mL) was added at 8 hours after transfection to trigger target gene expression.

